# Modeling of transcriptomic variation among subgenomes in 25 accessions of common wheat reveals cis- and trans- regulation architectures

**DOI:** 10.1101/2024.10.09.617331

**Authors:** Yasuyuki Nomura, Moeko Okada, Toshiaki Tameshige, Shotaro Takenaka, Kentaro K. Shimizu, Shuhei Nasuda, Atsushi J. Nagano

**Affiliations:** Research Institute for Food and Agriculture, Ryukoku University, 1-5 Yokotani, Seta Oe-cho, Otsu, Shiga 520-2194, Japan; Kihara Institute for Biological Research, Yokohama City University, 641-12 Maioka-cho, Totsuka-ku, Yokohama, Kanagawa 244-0813, Japan; Department of Evolutionary Biology and Environmental Studies, University of Zurich, Winterthurerstrasse 190-8057, Zurich; Graduate School of Science and Technology, Niigata University, 8050 Ikarashi 2-no-cho, Nishi-ku, Niigata, 950-2181, Japan; Graduate School of Life and Environmental Sciences, Kyoto Prefectural University, 1-5 Nakaragi-cho, Shimogamo, Sakyo-ku, Kyoto 606-8522 Japan; Faculty of Agriculture, Ryukoku University, 1-5 Yokotani, Seta Oe-cho, Otsu, Shiga 520-2194, Japan; Graduate school of Agriculture, Kyoto University, Kitashirakawaoiwake-cho, Sakyo-ku, Kyoto 606-8502, Japan; Institute for Advanced Biosciences, Keio University, 246-2, Tsuruoka, Yamagata 997-0052, Japan

**Keywords:** common wheat, transcriptome, Lasy-Seq, allohexaploid, *cis*/*trans*-regulation

## Abstract

Common wheat is an allohexaploid plant, thus making it difficult to obtain homoeolog-distinguished transcriptome data. Lasy-Seq, a type of 3’ RNA-seq, is efficient for obtaining homoeolog-distinguished transcriptomes and can thus overcome this measurement difficulty. This study obtained transcriptome data from the seedlings, second leaves, and root tips of 25 lines from mainly eastern transmitted area using Lasy-Seq. Roots and seedlings exhibited similar transcriptome profiles; however, they were different from those of the leaves. We determined the effects of subgenomes, lines and their interactions with leaves, roots, and seedlings on the expression levels of each homoeolog triad. Of the 19,805 homoeolog triads, 50.9–55.4%, 24.2–29.5%, and 7.7–9.0% showed significant effects on their expression levels from subgenome, line, and interaction, respectively. 51–55% and 24–30% have genetic variation in the *cis*- and *trans*-regulation. Hierarchical clustering and co-*trans* regulation network analysis of homoeolog triads revealed that the patterns of expression polymorphisms among the lines were shared in different genes. The triads in which the statistical model detected as line effects imply that expression variation between lines is caused by changes in a smaller number of common *trans*-factors. We assigned gene ontology (GO) terms of the *Arabidopsis* orthologs to wheat homoeolog triads via reciprocal BLAST between common wheat and *Arabidopsis*, thus improving the percentage of gene-assigned GO terms to all analyzed GO terms from 19.1% to 90.6%. GO term enrichment analysis revealed that GO terms related to each tissue type function were enriched in genes expressed in the leaves and roots. Our information provides fundamental knowledge for the future breeding of plants possessing complex gene regulatory networks such as common wheat.

## INTRODUCTION

Common wheat (*Triticum aestivum*) is widely cultivated and is the major cereal that accounts for approximately 20% of calories consumed worldwide. Unlike other major crops such as rice (diploid with a genome size of approximately 400 Mb), wheat possesses a large genome size (17 Gb) and is allohexaploid (2*n* = 6*x* = 42, AABBDD) and derived from interspecific hybridization. The three gene copies that originated from the parental species are highly similar and are termed homoeologs. Despite the relatively complex genome of wheat, the complete genome (IWGSC 2014, 2018) and transcriptome references (Zhu et al. 2021) have been developed. Using this information, understanding the regulation of gene expression is essential for efficient wheat breeding. A major difference compared to diploid crops is that common wheat possesses a complex gene expression network due to the three subgenomes of allohexaploid species derived from hybridization and genome duplication. Allopolyploidization results in the combination of different genomes, and thus, expression patterns can be inherited and combined to confer broad environmental tolerance (Stebbins 1971, Shimizu 2022). Furthermore, this can lead to new interactions between genomes (Hu and Wendel 2019) and changes in gene expression that have not yet been observed in the parental species (Yoo et al. 2013, Quan et al. 2022).

Expression quantitative trait locus (eQTL) analysis and expression genome-wide association study (eGWAS), in which gene expression levels are used as phenotypes, are useful tools for searching for loci that correlate with variations in gene expression (Gan et al. 2011, Zan et al. 2016, Wang et al. 2021). *Cis-*eQTLs are loci on the same chromosomes as that of the expressed gene and are also located near the expressed gene (often assumed to be within 1 Mb) (Zan et al. 2016). In contrast, *trans-*eQTLs are loci on different chromosomes from those carrying the expressed genes. Allohexaploid plants such as wheat can be used to distinguish *cis-* and *trans-*eQTLs without forward genetic mapping of the gene expression levels. Gene expression bias (homoeolog expression bias) among the subgenomes was most likely caused by *cis*-eQTLs that are unique to the subgenome. When there are differences in the average expression levels of all genomes compared to that of multiple lines, *trans*-eQTLs are likely responsible. Hence, wheat is a powerful tool for detecting homoeolog-specific expression and gene expression regulation due to *cis*- and *trans*-eQTLs (Wang et al. 2006, Shi et al. 2012, Yoo et al. 2013, Paape et al. 2016).

Genome-wide measurement of gene expression-level polymorphisms in wheat is difficult due to similarities among homoeolog sequences. Lasy-Seq, a type of 3’ RNA-seq, is efficient for obtaining homoeolog-distinguished transcriptomes (Kamitani et al. 2019, Sun et al. 2023) and can thus overcome this measurement difficulty. Lasy-Seq is a method for preparing libraries using oligo dT primers containing index sequences that allow for the mixing of samples from a relatively early stage of library preparation, thus reducing technical effort and experimental costs.

Common wheat originated from the Middle East (Dvorak et al. 2012) and spread westward to Europe and eastward to Asia. A study examining the genetic variation in common wheat revealed that landraces and cultivars from Asia diverged from those from Europe and underwent considerable variation (Takenaka et al. 2018). Recently, the chromosomal-level assembly of a representative Japanese cultivar, Norin 61, as well as that of an experimental strain, Chinese Spring, was reported (Walkowiak et al. 2020, Tsujimoto 2021, Shimizu et al. 2021). Although Asian cultivars have been underexploited in the context of modern breeding (Balfourier et al. 2019), many agriculturally relevant loci, such as semi-dwarfism, introgressed to facilitate the green revolution originating from the Asian germplasm, and this is the underlying premise for the uniqueness of the Asian germplasm.

In this study, we used 25 wheat lines from the eastern transmission area (Takenaka et al. 2018) provided by the National BioResource Project (NBRP) KOMUGI (https://shigen.nig.ac.jp/wheat/komugi/strains/aboutNbrp.jsp) in Japan. Lasy-Seq was used to obtain the leaf, root, and seedling transcriptomes of the lines. Based on these data, we detected gene expression regulation due to *cis-* or *trans-*eQTLs with respect to leaves, roots, and seedlings in wheats.

## MATERIALS AND METHODS

### Plant materials and common settings of the growth chamber

For stratification, 8 seeds from 25 wheat lines (Takenaka et al. 2018) were sown on filter paper in plastic petri dishes (5 cm diameter) and maintained in the dark at 4°C for 3 days. After stratification, half of seeds were maintained at 20°C with 24 h light for 1 day. Whole seedlings (endosperms, coleoptiles, and roots) of approximately 2-3 mm in size were sampled and then immediately frozen in liquid nitrogen and stored at -80°C (Figure 1A, B).

**Figure 1.**
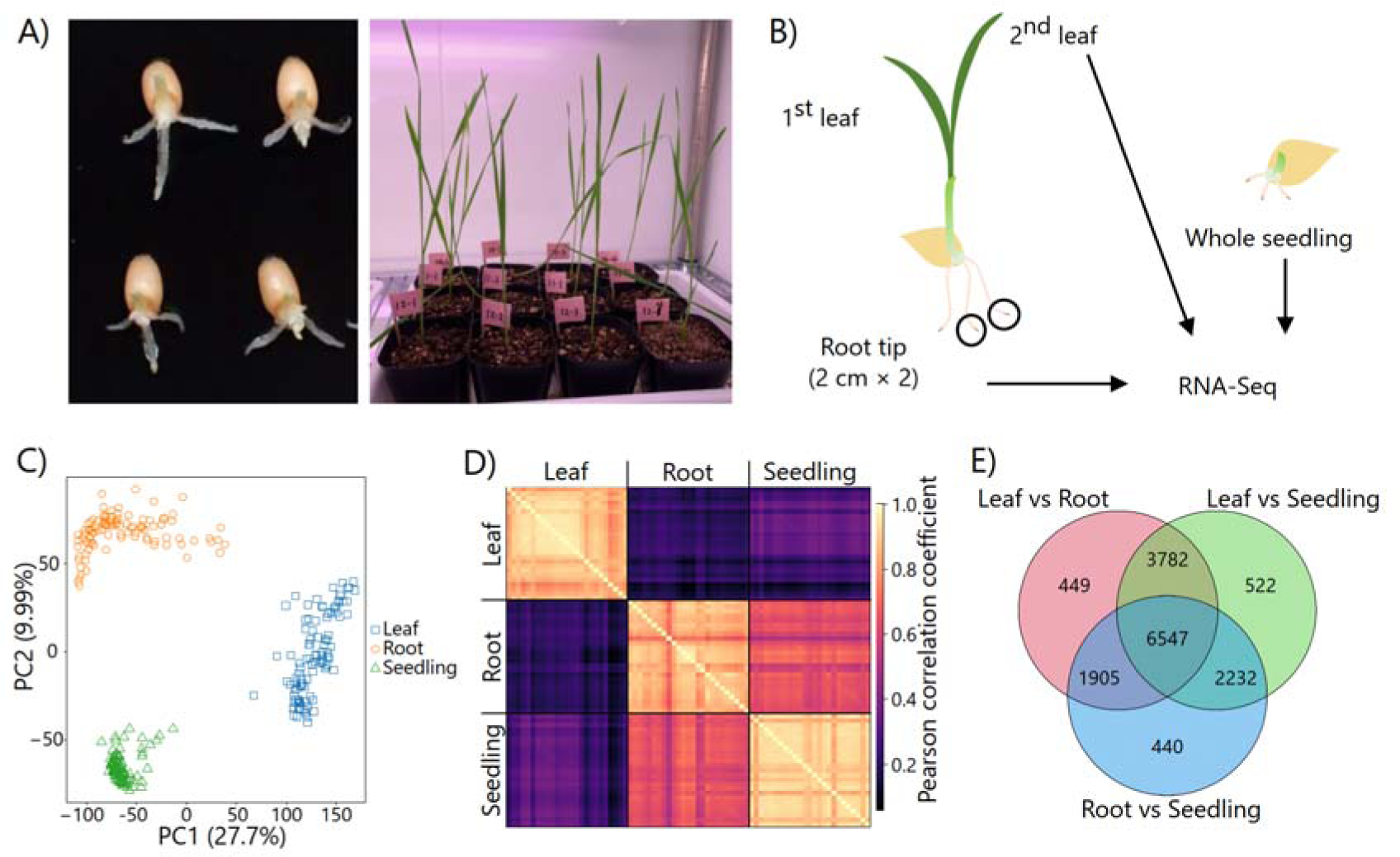
Summary of transcriptome profile in leaves, roots and seedlings. (A) Plants in sampling timing. (B) Sampling position in plants. (C) PCA of expressed genes (log_2_(RPM+1) > 2) for all samples. (D) Pearson correlation between the average of expressed genes by lines in leaves, roots and seedlings. The color bar on right side represents Pearson correlation coefficients. (E) Number of DEGs between tissue types.

Photosynthetic photon flux density (PPFD) was set at approximately 74 μ mol s^−1^ m^−2^ at the petri dish surface.

The rest of the seeds were sown into pots (7 cm diameter) that were filled with vermiculite absorbing liquid fertilizer (N; 1.5 mg, P; 2.5 mg, K; 1.3 mg per pot, HYPONeX, HYPONeX Japan, Osaka, Japan), and the pots were then maintained at 20°C with 24 h light. After 7–11 days, we sampled fully expanded second leaves and 2 cm of two fresh root tips (Figure 1A, B). These samples were immediately frozen in liquid nitrogen and stored at -80°C. PPFD was set at approximately 106 μ mol s^−1^ m^−2^ at the soil surface.

### RNA extraction, library preparation, and RNA-Seq

Leaves, roots, and seedlings (tissue type) were crushed with three zirconia beads using the TissueLyser II (QIAGEN, MD, USA) with pre-chilled adapters at -80 °C. Total RNA was extracted from the leaves using a Maxwell 16 LEV Plant RNA Kit (Promega, WI, USA). The amount of RNA was quantified using the Quant-iT RNA Assay Kit broad range (Thermo Fisher Scientific, Waltham, MA, USA) and a Tecan Infinite 200 PRO plate reader (Tecan, Männedorf, Switzerland). For the RNA-Seq library preparation, 400 or 500 ng of total RNA per sample was used. Libraries were prepared using the Lasy-Seq v1.1 protocol (Kamitani et al. 2019) (https://sites.google.com/view/lasy-seq/). Library quality was assessed using a Bioanalyzer (Agilent Technologies, Japan). Libraries were sequenced using paired-end sequencing (150 bases) on a HiSeq X (Illumina).

### Identification of homoeologous regions and homoeolog triads

The homoeolog triad lists based on IWGSC RefSeq v1.1 (IWGSC 2018) and v2.1 (Zhu et al. 2021) were prepared according to a previous study (Kuo et al. 2020, Sun et al. 2023). The transcript sequences of both high- and low-confidence genes from subgenomes A, B, and D were extracted based on the IWGSC reference sequences v1.0 and v2.0 and annotation information v1.1 and v2.1 of Chinese Spring. Homoeolog pairs within the AB, AD, and BD subgenomes were established by identifying the reciprocal best hits between transcripts from one and the other two subgenomes using LAST v950 (Kiełbasa et al. 2011). Homoeologous triads for both v1.1 and v2.1 were determined as homoeologs within the AB subgenomes that also exhibited identical hits for the D subgenome in their AD and BD pairs. Corresponding homoeologous triads between v1.1 and 2.1 were defined based on the reciprocal best hit of the A subgenome transcripts of each homoeologous triad.

### Preprocessing and handling of RNA-Seq data

Illumina adapter removal and low-quality filtering (Q < 20) of the RNA-Seq data were performed using fastp (Chen et al. 2018). We prepared the transcriptome reference by extending 1 kb on the 3’-end of transcriptome sequences in IWGSC RefSeq v2.1 (Zhu et al. 2021). Preprocessed reads were mapped onto the reference sequences using Bowtie (v1.1.1) (Langmead et al. 2009) and quantified using RSEM-1.2.21 (Li and Dewey 2011). Based on homoeolog sequence polymorphisms, RSEM was expected to assign read counts among the homoeologs. The conversion of the output from RSEM to reads per million (RPM) was performed using R (version 4) (R Core Team 2021) in the same manner as that used in a previous study (Kamitani et al. 2016). The read counts of all the transcript variants were summed to obtain the locus count. Based on a previous study (Sun et al. 2023), 19,856 homoeolog triads with known correspondence among the A, B, and D genomes in RefSeq v1.2 were included in the analysis. Of these, we analyzed 19,805 triads that corresponded to RefSeq v1.2 and v2.1.

To examine the accuracy of mapping distinguishing homoeologs by RSEM, we further mapped the RNA-Seq data of six Tetra Chinese Spring (TCS) lines that were obtained in a previous study (Sun et al. 2023) to RefSeq. The TCS subgenome D was removed from Chinese Spring by repeated backcrossing (Yang et al. 1999). The percentage of TCS read counts that map to the D genome is considered the error rate of mapping between homoeologs.

### Statistical analysis for RNA-Seq data

Principal component analysis (PCA) was conducted using the log_2_ (RPM+1) of the expressed genes (average log_2_ (RPM+1) > 2) in the homoeolog triads (Supplementary Figure S1). Pairwise Pearson’s correlations were calculated using gene expression averaged by line.

Differentially expressed genes (DEGs) and homoeolog triads (DETs) were detected using the edgeR package (Robinson et al. 2010). A model was constructed with three tissue types (leaves, roots and seedlings) as explanatory variables to detect DEGs among the tissue types. A model was constructed with 25 lines, three subgenomes (A, B, and D), and their interactions in each tissue type to detect DETs. A likelihood ratio test (LRT) was used to assess the significance level of the fixed effects. *P*-values adjusted using Holm’s method (Holm 1979) for the interaction term were calculated using LRT between models with and without the interaction term. Adjusted *P*-values for the line and genome terms were calculated by comparing the model without the interaction term to the model without the interaction and the line or subgenome terms. DEG detection was conducted with an adjusted *P*-value < 0.05.

Multiple regression quadratic assignment procedures (MRQAP) tests (Dekker et al. 2007) were performed to examine the effects of total read counts and genetic distance on the transcriptome Pearson’s correlation among 25 lines in each tissue type using “asnipe” package (Farine 2013). The geometric mean of the total read counts was used as an explanatory variable. Genetic distances were calculated using the Tajima-Nei model in MEGA11 (Tamura et al. 2021) based on 9,746 SNPs (Takenaka et al. 2018). Wilcoxon rank-sum tests were performed on the adjusted *P*-values of the line and subgenome terms between DEGs and non-DEGs between leaves and roots.

A weighted gene co-expression network analysis (WGCNA) was conducted for DETs for line terms to find candidate genes as *trans*-factor. We selected DETs whose variance of the mean expression, logarithmic coefficient, per line calculated by edgeR was greater than 1. DETs with a Spearman’s correlation coefficient of 0.8 or higher among the DETs were selected. WGCNA based on TOM similarity and Hierarchical clustering were performed using scaled logarithmic coefficient of the DETs. We used “WGCNA” package(Langfelder and Horvath 2012) to perform WGCNA. Only modules containing 10 or more DETs were used to draw the network.

*Preparation of GO annotations and enrichment analysis using the GO annotations* Reciprocal nucleotide BLAST between IWGSC RefSeq v2.1 (Zhu et al. 2021) and the *Arabidopsis thaliana* transcriptome reference (Araport11) (Cheng et al. 2017) was conducted to add gene ontology (GO) terms to the wheat transcriptome reference. GO terms were assigned to homoeologous triads. GO enrichment analysis was performed on the DEGs and DETs using Fisher’s exact test as described previously (Nagano et al. 2019). The *P*-values were adjusted by Holm’s method.

Hierarchical clustering was performed using GO terms that were enriched for significant genes in the subgenome, line, and interaction terms among expression profiles of the tissue types. For the four combinations (two transcriptome profiles [leaves and roots] × two terms in the model [subgenome and line]), the input data were generated as 1 if the GO terms were significantly enriched and as 0 if they were not.

## RESULTS

### New GO annotation list allowed for enrichment analysis

We prepared GO annotations of wheat based on *A. thaliana* GO annotations, as the references provided by the IWGSC did not provide enough GO terms to allow for the analysis to be performed. There were a small number of GO terms assigned to wheat genes (19.1% of the 10,334 terms) in the original reference (Zhu et al. 2021). We conducted a reciprocal nucleotide BLAST between wheat and *A. thaliana* to add GO terms to the wheat reference, and 40.6% of the 295,922 genes were reciprocal or one-sided hits to *A. thaliana* reference, ultimately resulting in numerous GO terms being assigned to at least one gene (90.6% of the 10,334 terms). As predicted based on the allohexaploid nature of wheat, approximately three-fold more *A. thaliana* genes were assigned per GO term (Supplementary Figure S2, Supplementary Table S1).

*Leaves exhibit different expression profiles compared to those of seedlings and roots* Approximately 4.4% of the TCS (Tetra Chinese Spring) reads were mapped to the D genome by Bowtie and RSEM (Supplementary Table S2). The results indicated that RSEM can be used for mapping to distinguish homoeologs. RNA-Seq was performed on three tissue types (seedlings, leaves and roots) from 25 wheat lines (Figure 1A, B, Supplementary Table S3). The average read counts for seedlings, leaves, and roots were 2.16±1.00, 2.49±1.83 and 2.47±2.34 M reads per sample, respectively. The mapped reads for seedlings, leaves, and roots in transcriptome reference were on average 71.2±3.9, 60.4±10.4, and 62.3±6.8%, respectively. The mapped reads for homoeolog triads of seedlings, leaves, and roots were 63.7±2.3, 51.3±9.9, and 39.9±17.2% (mean ±s.d.), respectively, and this indicated larger variation in roots than that in other tissue types. Root samples exhibited more reads mapped to rRNAs. Possibly, this is due to soil-derived contaminants that reduce stringency during the reverse transcription of root samples.

Of the 19,805 homoeolog triads (19,805 triads × 3 subgenomes = 59,415 genes), 32,021 genes were expressed (Supplementary Figure S1). PCA summarizing the gene expression profiles of the three tissue types was conducted (Figure 1C), and the results indicated that the PC1 and PC2 explained 27.7% and 10.0% of the variation, respectively. The gene expression profiles were grouped into three clusters corresponding to the tissue type, with the seedling profile being closer to the root profile. This was also supported by Pearson’s correlation analysis (Figure 1D). The geometric mean of the total read counts significantly affected the transcriptome correlation in the all tissue types by MRQAP tests (permutation *P*-value = 0.0, 2.7×10^-3^ and 7.4×10^-3^ in leaves, roots and seedlings, respectively) (Supplementary Table S4). Genetic distance also significantly affected the leaf and seedling correlation (*P*-value = 0.0178, 0.103 and 0.0343 in leaves, roots and seedlings, respectively).

In Chinese Spring, 2,262 DETs (differentially expressed homoeolog triads) between leaves and roots were shared with those between leaves and seedlings. GO enrichment analysis of the 2,262 DETs revealed that 140 GO terms, including plastid- and photosynthesis-related terms, were significantly enriched (Supplementary Table S5). Similar results were obtained when 25 lines were pooled for this analysis (Figure 1E, Supplementary Table S6).

### Analysis using subgenomes and lines as explanatory variables for DETs reveals cis- and trans-regulation

We analyzed the effects of the subgenomes and lines on gene expression. The gene expression level of a sample can be explained by the subgenome, the line to which the gene belongs, and their interactions (Figure 2A). The *cis*-element is expected to affect gene expression near the gene, but it does not affect gene expression in different subgenomes. Thus, DETs among subgenomes were expected to include the effects of factors that affect only certain subgenomes, primarily *cis*-elements (Figure 2A). Conversely, the *trans*-factors would not necessarily be located near the affected gene but would spread throughout the nucleus and affect gene expression in all subgenomes. Thus, DETs among lines are expected to primarily include the effects of *trans*-factors, as it is unlikely that a particular line possesses *cis*-mutations in all subgenomes. These effects occurred simultaneously. Differences in gene expression between subgenomes were shared across lines but the average gene expression levels differed among lines. In such cases, the statistical model detected both significant effects and no interaction terms (Supplementary Figure S3). If an interaction is detected, various gene-regulatory mechanisms are possible. For example, when a *cis*-element is present in particular subgenomes of particular lines (Supplementary Figure S3A) and when a *cis*- and *trans*-factor interact, this is indicative of gene-regulatory mechanisms (Supplementary Figure S3B, C).

**Figure 2.**
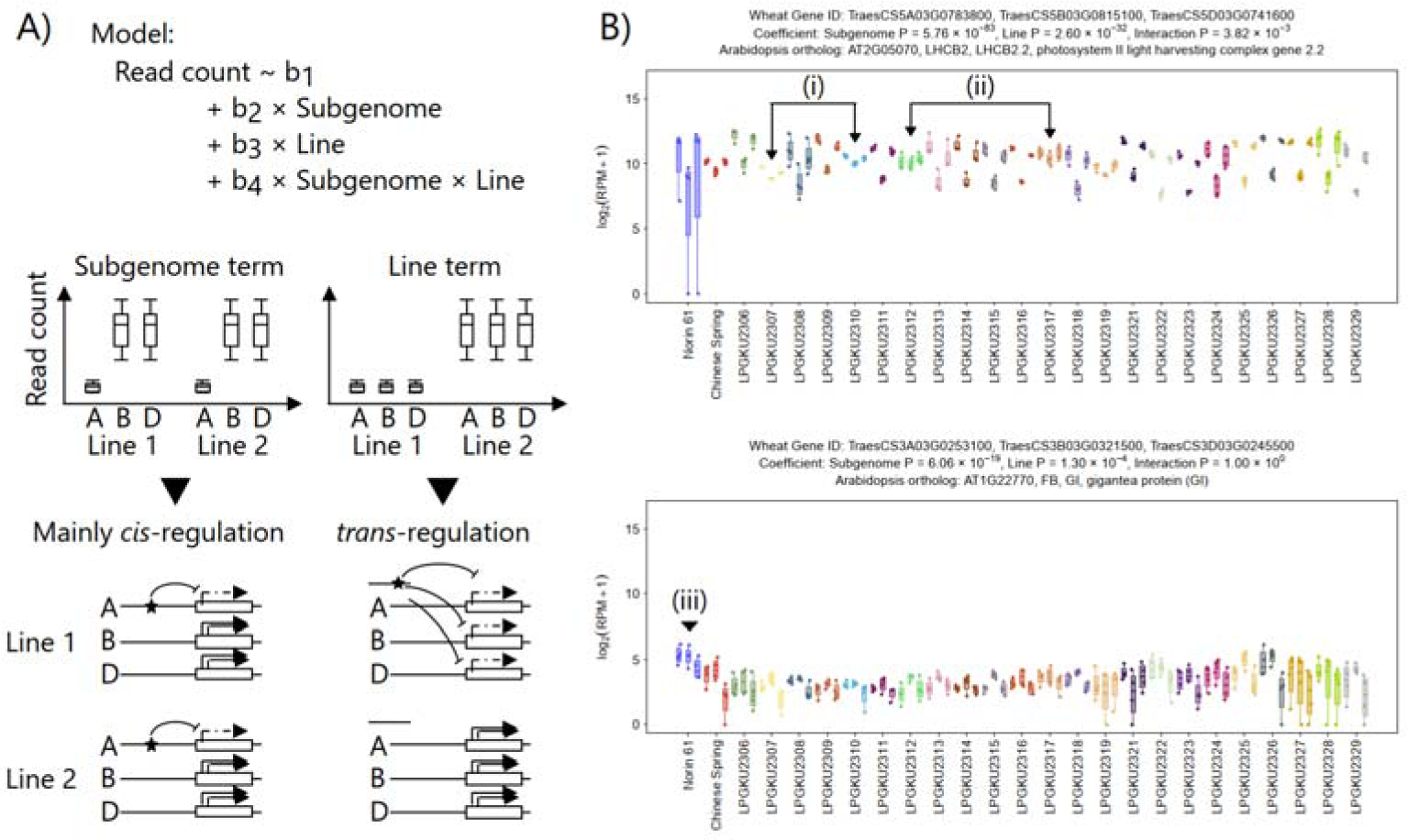
Relationship between statistics model and cis/trans-regulation. (A) Gene regulations estimated by significant terms in statistical models. (B) The expression level of *LHCB1* and *GI* in leaf. (i) represents a comparison of accessions with similar ratios of expression among subgenomes. (ii) represents accessions with lacking expression difference among subgenomes. (iii) represents *GI* expression of Norin 61.

Here, we present two examples that include *LHCB2.2* and *GIGANTEA* (*GI*). An example of a gene related to photosynthesis is the gene expression of *LHCB2.2* in leaves (Figure 2B), where *LHCB2.2* exhibited significantly different expression in subgenomes, lines, and their interactions (adjusted *P*-value = 5.76×10^-83^, 2.60×10^-32^ and 3.82×10^-3^, respectively). The expression ratio of subgenome B was lower than that of subgenomes A and D in all but two lines (LPGKU2312 and LPGKU2317, (ii) in Figure 2B), thus suggesting that *cis*-elements that suppress subgenome B expression are common in many lines. When comparing LPGKU2307 and LPGKU2310, the expression ratio between these subgenomes was similar, but *LHCB2.2* is less expressed in LPGKU2307 than it is in LPGKU2310 ((i) in Figure 2B). The variation of expression level is expected to occur via *trans*-factors in these lines.

*GI* is a circadian clock-controlled gene that promotes photoperiodic flowering in *Arabidopsis* and possibly in wheat under long-day conditions (Fowler et al. 1999, Xiang et al. 2005, Li et al. 2023) (Figure 2B). The expression of *GI* was significantly depended on the subgenome and line terms (adjusted *P*-value = 6.06×10^-19^ and 1.30×10^-4^, respectively). This gene was highly expressed in Norin 61 ((iii) in Figure 2B), which is consistent with the early flowering of Norin 61 (Shimizu et al. 2021).

Based on this idea, we examined the significance of the subgenome, line, and their interaction terms in the expression of 19,805 homoeolog triads. In seedlings, 55.4%, 24.2% and 9.0% were DETs for the subgenome, the line term and the interaction term, respectively (Figure 3A). For example, *Ribosomal protein S13/S15* (AT3G60770), *ATCATHB2* (AT1G02305) and *Ribosomal protein S3Ae* (AT4G34670) are significant only for subgenome, line and interaction term, respectively (Figure 3B, C). In leaves, 50.9%, 29.4% and 8.6% were DETs for the subgenome, the line term and the interaction term, respectively (Supplementary Figure S4). In roots, 52.4%, 27.3% and 7.7% were DETs for the subgenome, the line term and the interaction term, respectively (Supplementary Figure S5). In all tissue types, the number of triads expected to be *cis*-regulation was more than those expected to be *trans*-regulation.

**Figure 3.**
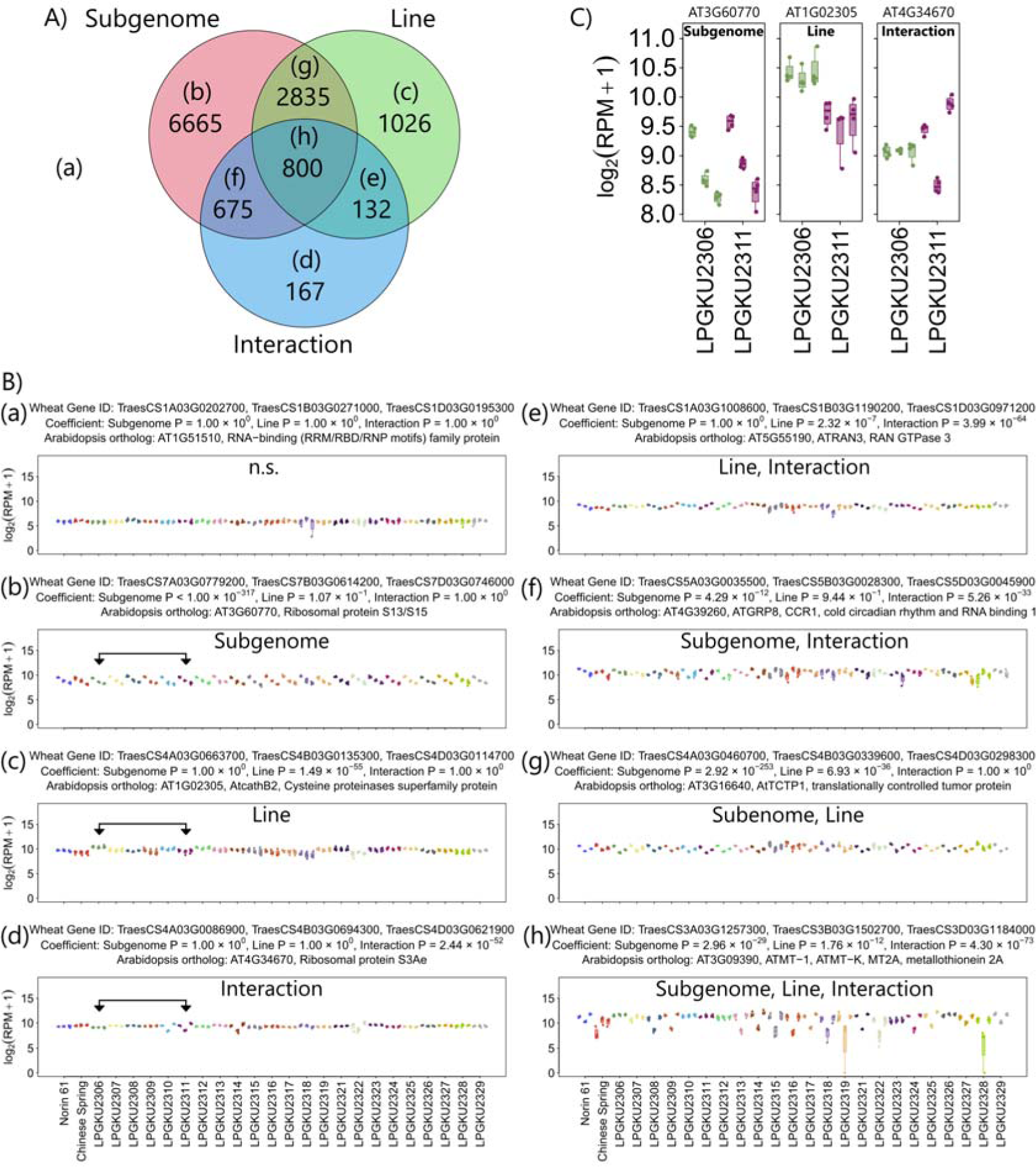
Expression levels of representative genes for which each term is significant in the seedlings. (A) The Venn diagram of the number of genes where line, subgenome or their interaction term is significant. (B) The representative genes corresponding to the Venn diagram; (a) all terms are not significant, (b) only subgenome term is significant, (c) only line term is significant, (d) only interaction term is significant, (e) line and interaction terms are significant, (f) subgenome and interaction terms are significant, (g) subgenome and line terms are significant, and (h) all terms are significant. Arrows represent LPGKU2306 and LPGKU2311. (C) The expression of *Ribosomal protein S13/S15*, *ATCATHB2* and *Ribosomal protein S3Ae* of LPGKU2306 and LPGKU2311.

### Functional characteristics of DETs

In the case of expressed genes in all tissue types (e.g., housekeeping genes), small differences in expression levels among lines were observed (Gan et al. 2011). Conversely, in the case of tissue-specific expressed genes that are not necessarily essential for fundamental cellular processes, differences in expression levels among lines are acceptable. Consistent with these predictions, DETs between leaves and roots exhibited significantly smaller adjusted *P*-values for subgenome and line terms for all tissue types compared to those of the non-DETs (Wilcoxon rank-sum test, *P* < 9.3 × 10^-53^; Supplementary Figure S6). This indicates that genes that are differentially expressed between leaves and roots (tissue type-specific expression) are more likely to be detected at different expression levels between lines and subgenomes.

In the transcriptome profiles of three tissue types, we assessed if the GO terms were enriched in genes with significant subgenome and line effects (Supplementary Table S7-9). A total of 315 GO terms were enriched in the three transcriptome profiles (Supplementary Table S10). The significant GO terms in the four combinations (two tissue types × two terms in the model) showed four patterns containing 10 or more GO terms (Figure 4). 87 GO terms including “RNA processing” (GO:0006396) were significantly enriched in line terms in leaves. 84 GO terms including “ribonucleoprotein complex” (GO:1990904) were significantly enriched in line terms in both leaves and roots. 49 GO terms including “response to stress” (GO:0006950) were significantly enriched in line terms in roots. 31 GO terms including “intercellular membrane-bounded organelle” (GO:0043231) were significantly enriched in both line and subgenome terms in leaves.

**Figure 4.**
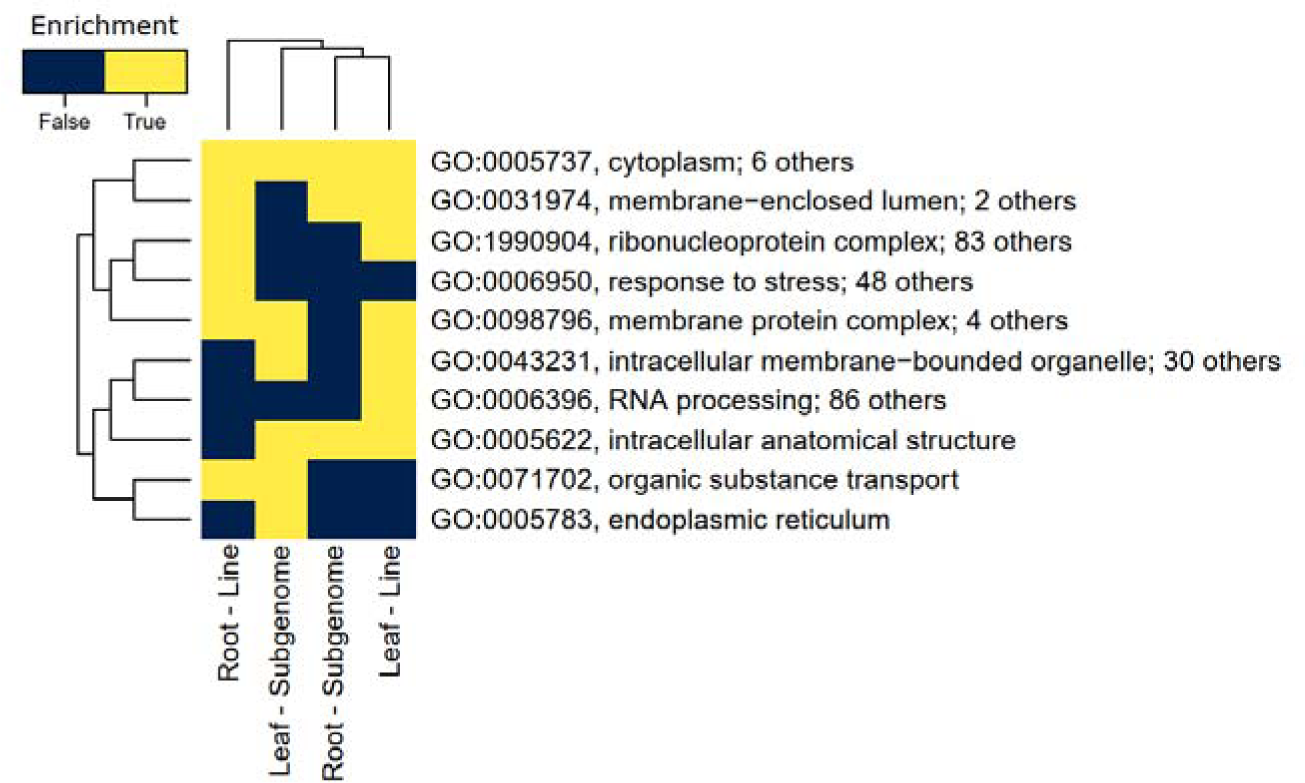
Significantly enriched GO terms in line and subgenome by leaves and roots. Heatmaps of GO terms. Dark and light colors represent non-enriched and enriched, respectively. Only one GO term of the same pattern is shown in row, see Supplementary Table S9 for a full list showing the rest. The dendrogram on the left and top of heatmap represents the result of hierarchical clustering. Distance in the dendrogram was based on Euclidean distance.

### Co-trans regulatory network analysis dissected trans-regulation architecture of transcriptome

For DETs for line terms, we conducted hierarchical clustering and WGCNA using the average triad expression levels, logarithmic coefficient, per line obtained from edgeR (Figure 5A). There were 5838 and 5403 DETs for line term for leaves and roots, respectively. Among them, there were 1247 and 985 DETs for which the variance of the logarithmic coefficient was greater than 1 for leaves and roots, respectively. Furthermore, there were 747 and 506 DETs with a spearman correlation coefficient of 0.8 or higher for leaves and roots, respectively. The triads expression patterns were divided into a few clusters in each tissue (Figure 5B). WGCNA based on TOM similarity showed that leaves were divided into three modules and roots into four modules (Figure 5C). The triads belonging to these different modules suggests that they are regulated by different *trans*-factors.

**Figure 5.**
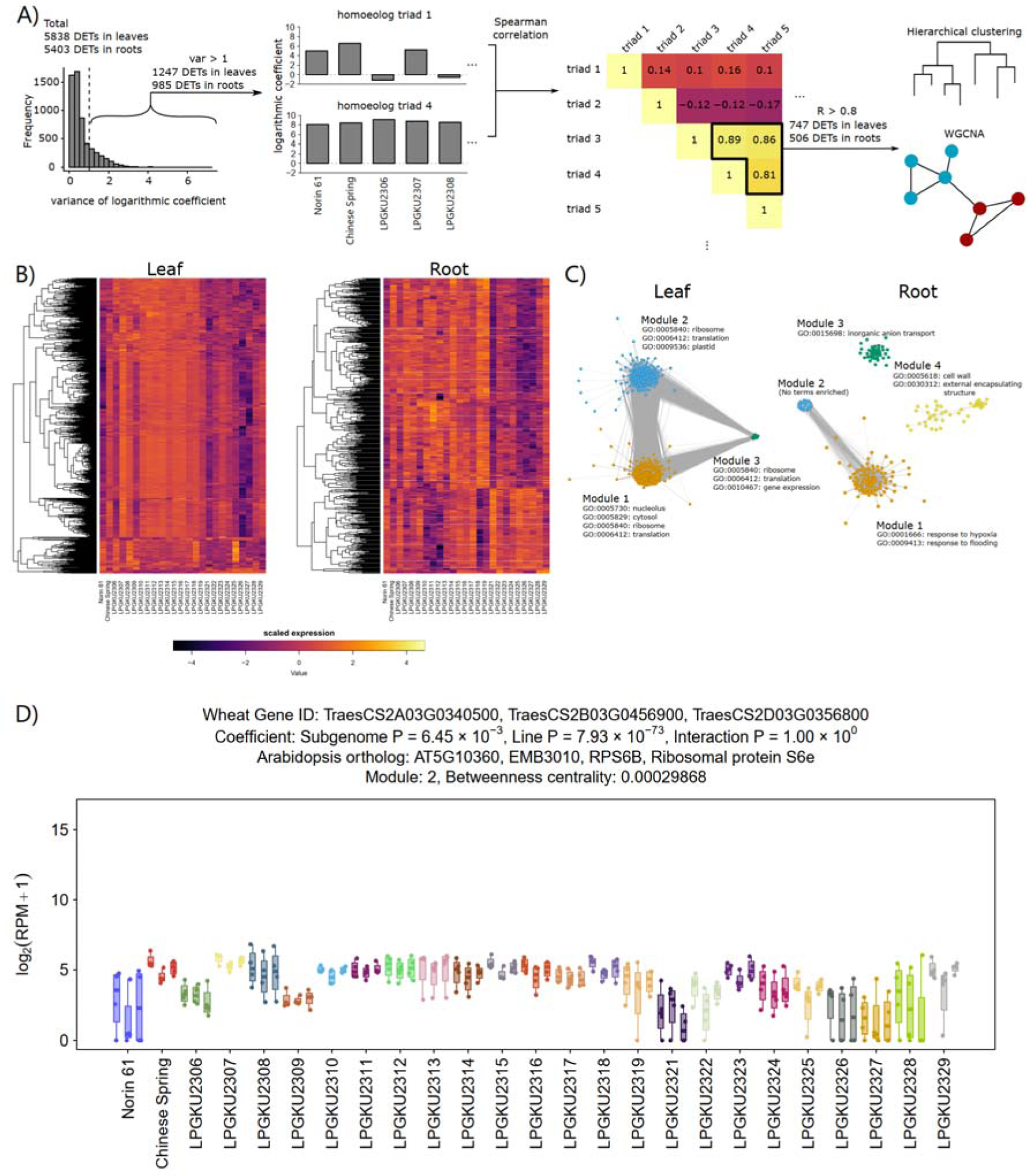
Hierarchical clustering and co-*trans* regulation network analysis of homoeolog triads. (A) Schematics of the analysis flow. For the mean expression per line (logarithmic coefficients) estimated by edgeR, triads with variance greater than 1 across lines are selected. The triads with Spearman correlations greater than 0.8 were subjected to hierarchical clustering and WGCNA. (B) Heat map and hierarchical clustering. Distance in the dendrogram was based on Euclidean distance. (C) WGCNA by TOM similarity. Some of the significant GO terms included in each module are shown. (D) The expression level in leaf of *RPS6B* which is a gene suspected to be a transcriptional factor of ribosomal proteins and with high betweenness centrality.

In leaves, ribosomal proteins were abundant in all modules and their betweenness centralities were high (Supplementary Table S11). GO enrichment analysis in each module showed that GO terms related to ribosomes was significantly enriched in all modules (Supplementary Table S12). One of the ribosomal proteins with high betweenness centrality is *RPS6B* which has been suggested in previous studies to act as a transcriptional regulator of rDNA and other ribosomal proteins (Kim et al. 2014). In roots, module 1 was characterized by *sucrose synthase 1* and *alcohol dehydrogenase 1* as triads with high betweenness centrality (Supplementary Table S13). Module 1 was enriched in GO in response to hypoxia and flooding (Supplementary Table S14). This may explain the differences in flooding tolerance that differed among the lines.

## DISCUSSION

In this study, the gene expression levels in the leaves, roots, and seedlings of 25 wheat lines were determined using Lasy-Seq (Kamitani et al. 2019) to distinguish homoeologs. Lasy-Seq can reduce experimental costs while maintaining high accuracy in gene expression quantification in plants or animals (Kashima et al. 2021, Yagi et al. 2021), but little analysis has been performed for allopolyploid plants. Here, we quantified the expression of different genes among the tissue types (Figure 1) and homoeologs (Figures 2, 3), thus demonstrating the utility of the Lasy-Seq method in the context of allohexaploids (Sun et al. 2023).

Norin 61 shows early flowering (Shimizu et al. 2021). In this study, we found high *GI* gene expression in Norin 61 (Figure 2). Because *GI* is suggested to be associated with early flowering in *Arabidopsis thaliana* (Fowler et al. 1999) and wheat (Xiang et al. 2005, Li et al. 2023), the early flowering of Norin 61 may be regulated by *GI*. As flowering time is a major breeding target for wheat, *GI* is a candidate gene that could be utilized for future breeding. Useful crops include a large number of allopolyploid plants such as kiwi fruits, bananas, and potatoes (Akagi et al. 2022). The availability of low-cost transcriptome analysis will also allow us to understand the gene expression regulatory networks and origins of cultivation related to allopolyploid crops.

The regulatory mechanisms of transcription can be distinguished as *cis-* and *trans-*regulation. Previous studies, including eQTL analyses, have inferred *cis-*mutations (Keurentjes et al. 2007, Wittkopp and Kalay 2012, Zan et al. 2016) or *cis-* and *trans-*regulation from changes in gene expression levels between the parental and hybrid lines (Shi et al. 2012, Quan et al. 2022). We hypothesized that distinguishing gene expression from line and subgenome effects would aid in the understanding of transcriptional regulation (Figure 2). eQTL analysis indicated that *cis*-eQTLs are more common than *trans*-eQTLs (Li et al. 2006, Keurentjes et al. 2007, Zan et al. 2016) (but see also Diaz-Valenzuela *et al*. [2020]). Of the 19,805 homoeolog triads examined in this study, 51–55% and 24–30% were under *cis*- and *trans*-regulation, respectively (Figure 2C). In the native East Asian wheat population, similar to the results of previous studies, *cis*-regulation was still the primary cause of variation in expression levels. However, the observation that *trans*-regulation also accounts for a large proportion of expression variation indicates the importance of searching for *trans*-factors in eQTLs.

We assumed that significant subgenome effects likely indicated *cis*-regulatory variations. Here, we did not distinguish among the possible types of *cis*-effect variations (variation in promoter activity, mRNA stability and variation in gene dosage, etc.) that can alter the expression level of a single homoeolog. There were many single nucleotide polymorphisms (SNPs), present and absent variations (PAVs) or copy number variations (CNVs) between Chinese Spring and Norin 61 (Walkowiak et al. 2020, Shimizu et al. 2021) likely suggesting a large total number of SNP, PAV and CNV among the 25 lines used in this study. Further genomic studies of these lines would contribute to deeper insights into the molecular details of the evolution of the *cis*-regulatory system.

Hierarchical clustering and WGCNA analysis using coefficients of the model revealed that there are DETs that show similar expression patterns to each other among the DETs for line term (Figure 5B, C). The similar expression patterns strongly suggest the presence of *trans*-factors that regulate the expression of multiple triads. In the DETs in leaves, a large number of ribosome-associated genes showed high values of betweenness centrality (Figure 5C, Supplementary Table S11). *RPS6B*, included in ribosome-associated genes, is a component protein of the ribosome and involved in regulating the expression of rRNA and several ribosomal proteins (Kim et al. 2014). A number of ribosome-associated genes with similar expression is likely influenced by differences in *RPS6B* as a *trans*-factor.

Revealing a complete picture of the variations in *cis-* and *trans-*regulation of the transcriptome is challenging. In this study, by leveraging allohexaploid wheat lines (Takenaka et al. 2018) and high-throughput RNA-Seq (e.g., Lasy-Seq [Kamitani *et al*. 2019]), we identified genes with expression level polymorphisms regulated in *cis-* and/or *trans-* manners. To clarify the genes responsible for *trans*-regulation, an analysis of a large mapping population is required. Target RNA-Seq (Kashima et al. 2022) enables effective quantification of the expression of *trans*-regulated genes in thousands of lines. Future analyses using large mapping populations will reveal the regulation of each gene and its contribution to the yield.

## ACKNOWLEDGEMENTS

We greatly appreciate Fumie Kobayashi, Kyoko Mogami, Satoko Kondo and Yumi Suzuki for their help with RNA-Seq library preparation, Motohiro Mihara and Hitoshi Ooshima of Dynacom for their support in RNA-Seq data analysis, and Miyuki Nitta for providing germplasms. This study was supported by the Japan Science and Technology Agency (Grant numbers, JPMJCR16O3 to K.K.S., JPMJCR15O2 and JPMJFR210B to A.J.N.); Japan Society for the Promotion of Science (JP20H00423, JP23K18156 and JP23H00386 to A.J.N.); Japanese Ministry of Education, Culture, Sports, Science and Technology (JP22H05179 to K.K.S., and JP23H04967 to A.J.N.); Swiss National Science Foundation (31003A_182318 and 31003A_212551 to K.K.S.); University Research Priority Program “Global Change and Biodiversity” from the University of Zurich to K.K.S.; and Global Affairs of the University of Zurich and JSPS International Leading Research grant 22K21352 to K.K.S. and S.N.

## DATA AVAILABLITY

The RNA-Seq data were submitted to the NCBI Sequence Read Archive repository under the BioProject number PRJNA976071. R scripts and data used in the analysis (count data, etc.) are available via the Dryad and Zenodo repository (https://doi.org/10.5061/dryad.fqz612jzq).

## SUPPLEMENTARY MATERIAL

**Supplementary Figure S1 Summary of sample sets and gene sets used in this study.** The histogram of the total read numbers is provided in the inset. Gene sets analyzed in this study. The histogram of mean expressions is presented in the inset. The dashed line indicates the threshold of averaged log_2_(RPM+1) > 2.

**Supplementary Figure S2 The number of genes for each GO term in wheat and *A. thaliana*.** The left and right panels represent results using original IWGSC reference and our result from reciprocal BLAST, respectively. The two axes are log-transformed. The black and red lines indicate y = x and y = 3x, respectively.

**Supplementary Figure S3 Relationship between the statistics model with interaction term and *cis*/*trans*-regulation.** Gene regulation as estimated by significant terms in statistical models. Differences in gene expression between subgenomes are shared across lines, but the average gene expression levels differ between lines. In such cases, the statistical model would detect both effects significantly and indicate no interaction term. If interaction is detected, various mechanisms of gene regulations are possible. For example, a detected interaction may indicate that (A) a *cis*-element is present in particular subgenomes of particular lines or that (B, C) a *cis*- and *trans*-factor interact with each other.

**Supplementary Figure S4 Expression levels of representative genes for which each term is significant in the leaves.** The figure above shows the number of genes where line, subgenome or their interaction term is significant. It also shows the correspondence between the Venn diagram and representative genes; (a) all terms are not significant, (b) only the subgenome term is significant, (c) only the line term is significant, (d) only the interaction term is significant, (e) the line and interaction terms are significant, (f) the subgenome and interaction terms are significant, (g) the subgenome and line terms are significant, and (h) all terms are significant.

**Supplementary Figure S5 Expression levels of representative genes for which each term is significant in the roots.** The figure above shows the number of genes where line, subgenome or their interaction term is significant. It also shows the correspondence between the Venn diagram and representative genes; (a) all terms are not significant, (b) only the subgenome term is significant, (c) only the line term is significant, (d) only the interaction term is significant, (e) the line and interaction terms are significant, (f) the subgenome and interaction terms are significant, (g) the subgenome and line terms are significant, and (h) all terms are significant.

**Supplementary Figure S6 Distribution of adjusted *P*-value in the subgenome and line term.** The x-axis represents log-transformed adjusted *P*-value of the subgenome and line terms in all tissue types as calculated by edgeR. Wilcoxon rank-sum tests were performed to detect the differences in the distribution of adjusted *P*-values between DETs and non-DETs between leaves and roots.

**Supplementary Table S1 Newly prepared GO annotations of homoeolog triads of wheat**

**Supplementary Table S2 Read counts of TCS mapped to each genome**

**Supplementary Table S3 Sample list and index ID**

**Supplementary Table S4 The effects of total read counts and genetic distance to transcriptome correlations**

**Supplementary Table S5 Enriched GO terms in DEGs between leaves and others in Chinese Spring**

**Supplementary Table S6 Enriched GO terms in DEGs between leaves and others**

**Supplementary Table S7 Enriched GO terms in DETs in model of seedling**

**Supplementary Table S8 Enriched GO terms in DETs in model of leaves**

**Supplementary Table S9 Enriched GO terms in DETs in model of roots**

**Supplementary Table S10 Pattern of significantly enriched GO terms in terms in model by tissue types**

**Supplementary Table S11 Module, cluster and centrality of DETs in leaves**

**Supplementary Table S12 Enriched GO terms in DETs in modules in leaves**

**Supplementary Table S13 Module, cluster and centrality of DETs in roots**

**Supplementary Table S14 Enriched GO terms in DETs in modules in roots**

## Notes

https://doi.org/10.5061/dryad.fqz612jzq

### Competing Interest Statement

The authors have declared no competing interest.

## REFERENCES

Akagi, T., Jung, K., Masuda, K., and Shimizu, K.K. (2022) Polyploidy before and after domestication of crop species. Curr. Opin. Plant Biol. 69: 102255.

Balfourier, F., Bouchet, S., Robert, S., DeOliveira, R., Rimbert, H., Kitt, J., et al. (2019) Worldwide phylogeography and history of wheat genetic diversity. Sci. Adv. 5: eaav0536.

Chen, S., Zhou, Y., Chen, Y., and Gu, J. (2018) Fastp: An ultra-fast all-in-one FASTQ preprocessor. Bioinformatics 34: i884–i890.

Cheng, C.Y., Krishnakumar, V., Chan, A.P., Thibaud-Nissen, F., Schobel, S., and Town, C.D. (2017) Araport11: a complete reannotation of the *Arabidopsis thaliana* reference genome. Plant J. 89: 789–804.

Dekker, D., Krackhardt, D., and Snijders, T.A.B. (2007) Sensitivity of MRQAP tests to collinearity and autocorrelation conditions. Psychometrika 72: 563–581.

Diaz-Valenzuela, E., Sawers, R.H., and Cibrian-Jaramillo, A. (2020) *Cis*- and *trans*-regulatory variations in the domestication of the chili pepper fruit. Mol. Biol. Evol. 37: 1593–1603.

Dvorak, J., Deal, K.R., Luo, M.C., You, F.M., Von Borstel, K., and Dehghani, H. (2012) The origin of spelt and free-threshing hexaploid wheat. J. Hered. 103: 426–441.

Farine, D.R. (2013) Animal social network inference and permutations for ecologists in R using *asnipe*. Methods Ecol. Evol. 4: 1187–1194.

Fowler, S., Lee, K., Onouchi, H., Samach, A., Richardson, K., Morris, B., et al. (1999) *GIGANTEA*: A circadian clock-controlled gene that regulates photoperiodic flowering in *Arabidopsis* and encodes a protein with several possible membrane-spanning domains. EMBO J. 18: 4679–4688.

Gan, X., Stegle, O., Behr, J., Steffen, J.G., Drewe, P., Hildebrand, K.L., et al. (2011) Multiple reference genomes and transcriptomes for *Arabidopsis thaliana*. Nature 477: 419–423.

Holm, S. (1979) A simple sequentially rejective multiple test procedure. Scand. J. Stat. 6: 65–70.

Hu, G. and Wendel, J.F. (2019) *Cis–trans* controls and regulatory novelty accompanying allopolyploidization. New Phytol. 221: 1691–1700.

IWGSC. (2014) A chromosome-based draft sequence of the hexaploid bread wheat (*Triticum aestivum*) genome. Science (80-.). 345: 1250092.

IWGSC. (2018) Shifting the limits in wheat research and breeding using a fully annotated reference genome. Science (80-.). 361: eaar7191.

Kamitani, M., Kashima, M., Tezuka, A., and Nagano, A.J. (2019) Lasy-Seq: a high-throughput library preparation method for RNA-Seq and its application in the analysis of plant responses to fluctuating temperatures. Sci. Rep. 9: 7091.

Kamitani, M., Nagano, A.J., Honjo, M.N., and Kudoh, H. (2016) RNA-Seq reveals virus-virus and virus-plant interactions in nature. FEMS Microbiol. Ecol. 92: fiw176.

Kashima, M., Kamitani, M., Nomura, Y., Moriyama, N.M., Betsuyaku, S., Hirata, H., et al. (2022) DeLTaLSeqL: directLlysate targeted RNALSeq from crude tissue lysate. Plant Methods 18: 99.

Kashima, M., Shida, Y., Yamashiro, T., Hirata, H., and Kurosaka, H. (2021) Intracellular and intercellular gene regulatory network inference from time-course individual RNA-Seq. Front. Bioinforma. 1: 777299.

Keurentjes, J.J.B., Fu, J., Terpstra, I.R., Garcia, J.M., van den Ackerveken, G., Snoek, L.B., et al. (2007) Regulatory network construction in Arabidopsis by using genome-wide gene expression quantitative trait loci. Proc. Natl. Acad. Sci. U. S. A. 104: 1708–1713.

Kiełbasa, S.M., Wan, R., Sato, K., Horton, P., and Frith, M.C. (2011) Adaptive seeds tame genomic sequence comparison. Genome Res. 21: 487–493.

Kim, Y.K., Kim, S., Shin, Y.J., Hur, Y.S., Kim, W.Y., Lee, M.S., et al. (2014) Ribosomal protein S6, a target of rapamycin, is involved in the regulation of rRNA genes by possible epigenetic changes in *Arabidopsis*. J. Biol. Chem. 289: 3901–3912.

Kuo, T.C.Y.Y., Hatakeyama, M., Tameshige, T., Shimizu, K.K., and Sese, J. (2020) Homeolog expression quantification methods for allopolyploids. Brief. Bioinform. 21: 395–407.

Langfelder, P. and Horvath, S. (2012) Fast R functions for robust correlations and hierarchical clustering. J. Stat. Softw. 46: 1–17.

Langmead, B., Trapnell, C., Pop, M., and Salzberg, S.L. (2009) Ultrafast and memory-efficient alignment of short DNA sequences to the human genome. Genome Biol. 10: R25.

Li, B. and Dewey, C.N. (2011) RSEM: accurate transcript quantification from RNA-Seq data with or without a reference genome. BMC Bioinformatics 12: 323.

Li, C., Lin, H., Debernardi, J.M., and Zhang, C. (2023) *GIGANTEA* accelerates wheat heading time through gene interactions converging on *FLOWERING LOCUS T1*. bioRxiv. 10.1101/2023.07.11.548614

Li, Y., Álvarez, O.A., Gutteling, E.W., Tijsterman, M., Fu, J., Riksen, J.A.G., et al. (2006) Mapping determinants of gene expression plasticity by genetical genomics in *C. elegans*. PLoS Genet. 2: e222.

Nagano, A.J., Kawagoe, T., Sugisaka, J., Honjo, M.N., Iwayama, K., and Kudoh, H. (2019) Annual transcriptome dynamics in natural environments reveals plant seasonal adaptation. Nat. Plants 5: 74–83.

Paape, T., Hatakeyama, M., Shimizu-Inatsugi, R., Cereghetti, T., Onda, Y., Kenta, T., et al. (2016) Conserved but attenuated parental gene expression in allopolyploids: Constitutive zinc hyperaccumulation in the allotetraploid *Arabidopsis kamchatica*. Mol. Biol. Evol. 33: 2781–2800.

Quan, C., Chen, G., Li, S., Jia, Z., Yu, P., Tu, J., et al. (2022) Transcriptome shock in interspecific F1 allotriploid hybrids between *Brassica* species. J. Exp. Bot. 73: 2336–2353.

R Core Team. (2021) R: A language and environment for statistical computing. R A Lang. Environ. Stat. Comput. https://www.R-project.org/

Robinson, M.D., McCarthy, D.J., and Smyth, G.K. (2010) edgeR: A Bioconductor package for differential expression analysis of digital gene expression data. Bioinformatics 26: 139–140.

Shi, X., Ng, D.W.K., Zhang, C., Comai, L., Ye, W., and Chen, Z.J. (2012) *Cis-* and *trans-*regulatory divergence between progenitor species determines gene-expression novelty in *Arabidopsis* allopolyploids. Nat. Commun. 3: 950.

Shimizu, K.K. (2022) Robustness and the generalist niche of polyploid species: Genome shock or gradual evolution? Curr. Opin. Plant Biol. 69: 102292.

Shimizu, K.K., Copetti, D., Okada, M., Wicker, T., Tameshige, T., Hatakeyama, M., et al. (2021) De novo genome assembly of the Japanese wheat cultivar Norin 61 highlights functional variation in flowering time and *Fusarium*-resistant genes in East Asian genotypes. Plant Cell Physiol. 62: 8–27.

Stebbins, G.L. (1971) Chromosomal Evolution in Higher Plants. Edward Arnold Ltd.., London.

Sun, J., Okada, M., Tameshige, T., Shimizu-Inatsugi, R., Akiyama, R., Nagano, A.J., et al. (2023) A low-coverage 3’ RNA-seq to detect homeolog expression in polyploid wheat. NAR Genomics Bioinforma. 5: lqad067.

Takenaka, S., Nitta, M., and Nasuda, S. (2018) Population structure and association analyses of the core collection of hexaploid accessions conserved *ex situ* in the Japanese gene bank NBRP-Wheat. Genes Genet. Syst. 93: 237–254.

Tamura, K., Stecher, G., and Kumar, S. (2021) MEGA11: Molecular evolutionary genetics analysis version 11. Mol. Biol. Evol. 38: 3022–3027.

Tsujimoto, H. (2021) Gene-mining Asian wheat to feed the population in the 21st century. Plant Cell Physiol. 62: 1–2.

Walkowiak, S., Gao, L., Monat, C., Haberer, G., Kassa, M.T., Brinton, J., et al. (2020) Multiple wheat genomes reveal global variation in modern breeding. Nature 588: 277–283.

Wang, J., Tian, L., Lee, H.S., Wei, N.E., Jiang, H., Watson, B., et al. (2006) Genomewide nonadditive gene regulation in *Arabidopsis* allotetraploids. Genetics 172: 507–517.

Wang, Z., Yang, L., Wu, D., Zhang, N., and Hua, J. (2021) Polymorphisms in *cis*-elements confer *SAUR26* gene expression difference for thermo-response natural variation in Arabidopsis. New Phytol. 229: 2751–2764.

Wittkopp, P.J. and Kalay, G. (2012) *Cis*-regulatory elements: Molecular mechanisms and evolutionary processes underlying divergence. Nat. Rev. Genet. 13: 59–69.

Xiang, Y.Z., Mao, S.L., Jia, R.L., Chun, M.G., and Xian, S.Z. (2005) The wheat *TaGI1*, involved in photoperiodic flowering, encodes an Arabidopsis *GI* ortholog. Plant Mol. Biol. 58: 53–64.

Yagi, H., Nagano, A.J., Kim, J., Tamura, K., Mochizuki, N., Nagatani, A., et al. (2021) Fluorescent protein-based imaging and tissue-specific RNA-seq analysis of Arabidopsis hydathodes. J. Exp. Bot. 72: 1260–1270.

Yang, Y.F., Furuta, Y., Nagata, S.I., and Watanabe, N. (1999) Tetra Chinese spring with AABB genomes extracted from the hexaploid common wheat, Chinese Spring. Genes Genet. Syst. 74: 67–70.

Yoo, M.J., Szadkowski, E., and Wendel, J.F. (2013) Homoeolog expression bias and expression level dominance in allopolyploid cotton. Heredity (Edinb). 110: 171–180.

Zan, Y., Shen, X., Forsberg, S.K.G., and Carlborg, Ö. (2016) Genetic regulation of transcriptional variation in natural *Arabidopsis thaliana* accessions. G3 Genes, Genomes, Genet. 6: 2319–2328.

Zhu, T., Wang, L., Rimbert, H., Rodriguez, J.C., Deal, K.R., De Oliveira, R., et al. (2021) Optical maps refine the bread wheat *Triticum aestivum* cv. Chinese Spring genome assembly. Plant J. 107: 303–314.

