## Supplementary Figure for "Modeling of transcriptomic variation among subgenomes in 25 accessions of common wheat reveals cis- and trans- regulation architectures"

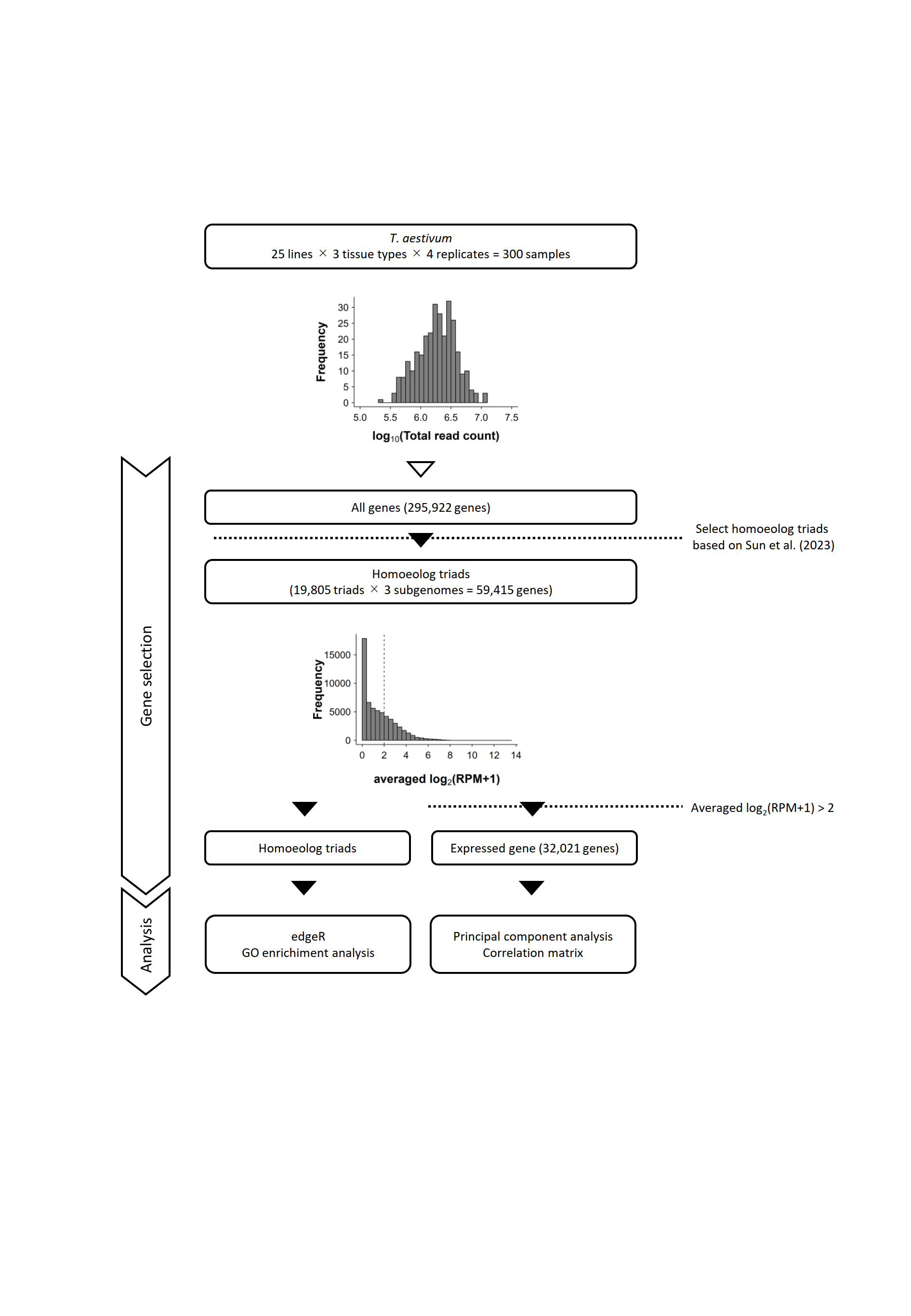

**Supplementary Figure S1 Summary of sample sets and gene sets used in this study.** The histogram of the total read numbers is provided in the inset. Gene sets analyzed in this study. The histogram of mean expressions is presented in the inset. The dashed line indicates the threshold of averaged log_2_(RPM+1) > 2.

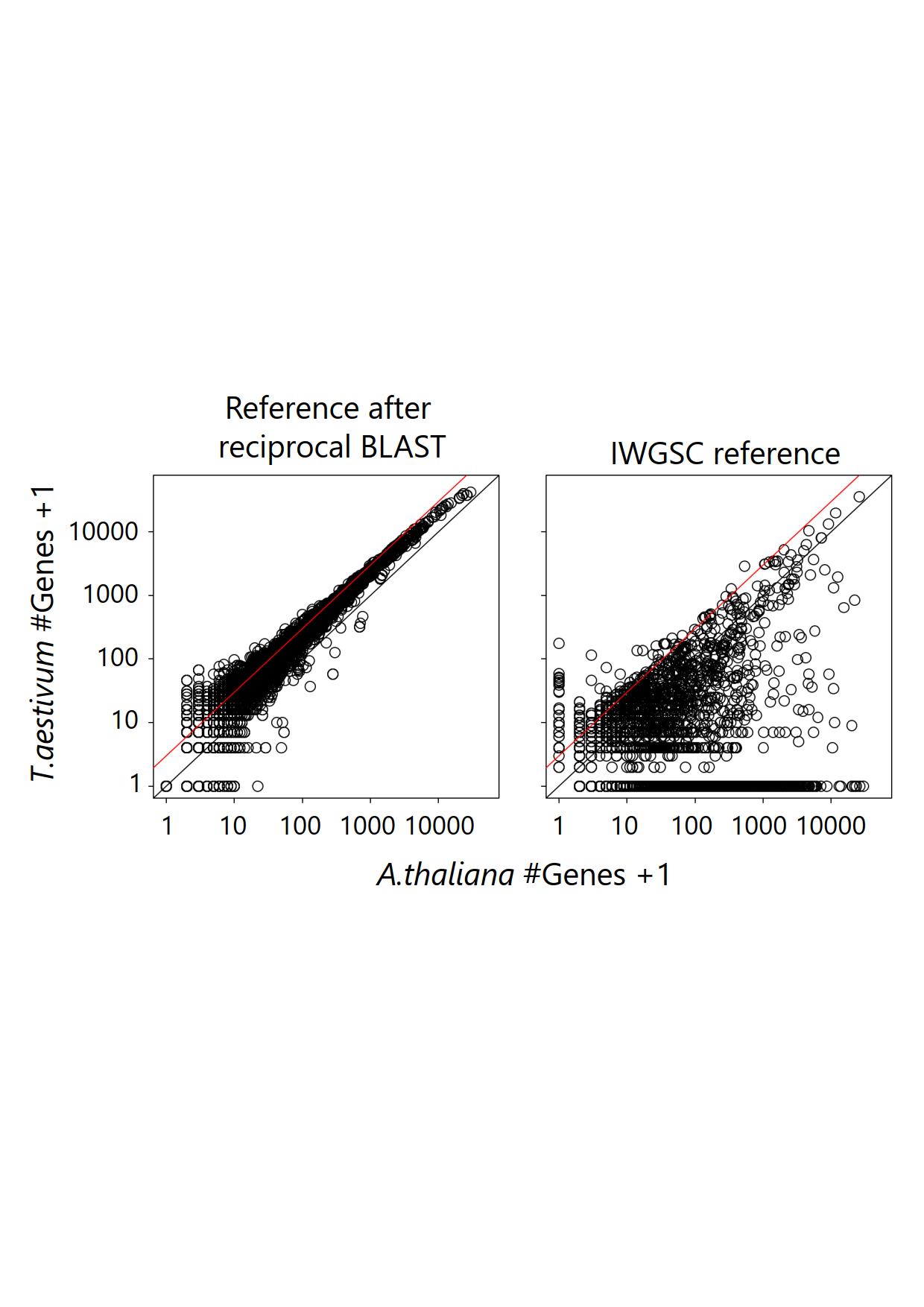

**Supplementary Figure S2 The number of genes for each GO term in wheat and *A. thaliana*.** The left and right panels represent results using original IWGSC reference and our result from reciprocal BLAST, respectively. The two axes are log-transformed. The black and red lines indicate y = x and y = 3x, respectively.

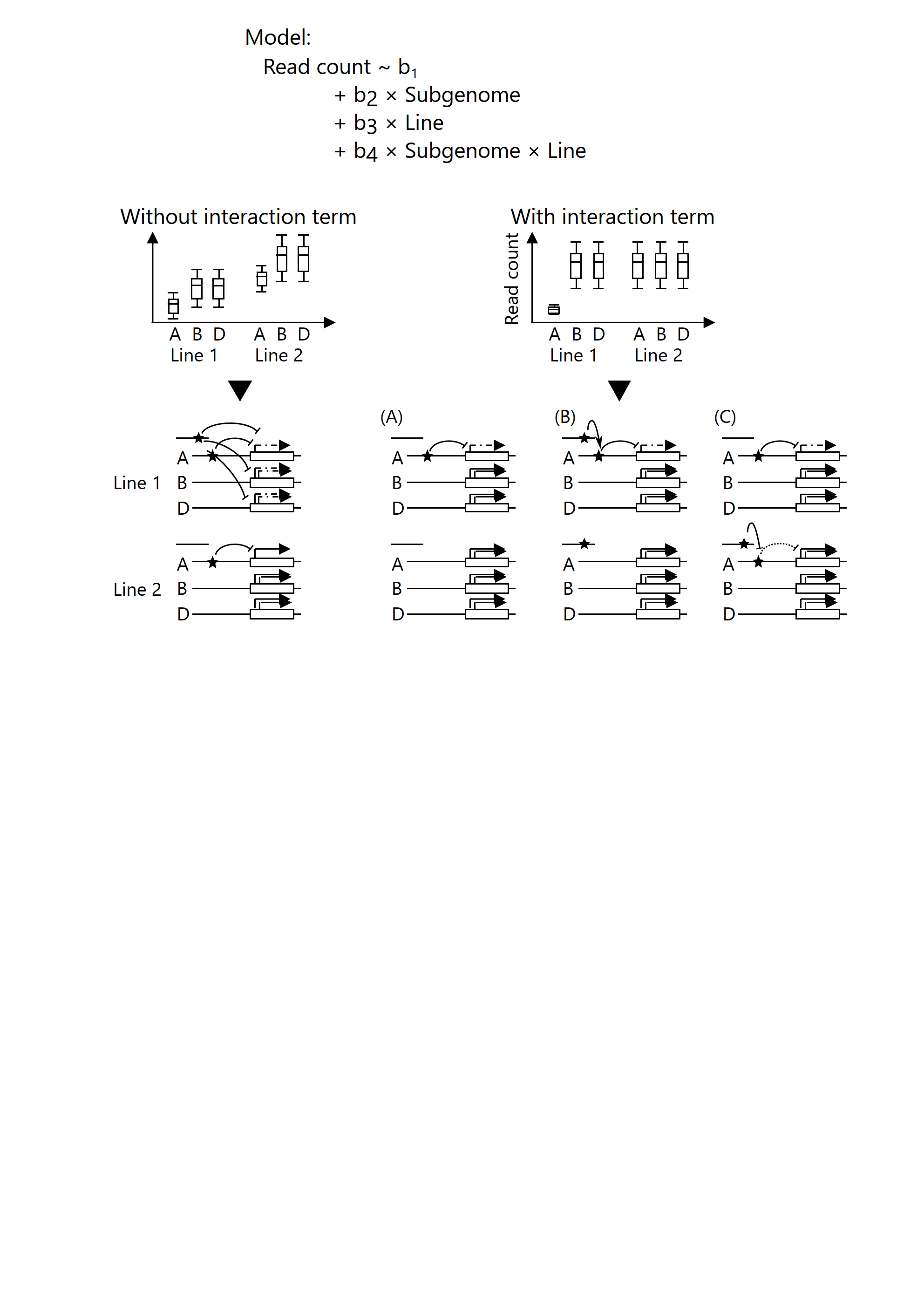

**Supplementary Figure S3 Relationship between the statistics model with interaction term and *cis*/*trans*-regulation.** Gene regulation as estimated by significant terms in statistical models. Differences in gene expression between subgenomes are shared across lines, but the average gene expression levels differ between lines. In such cases, the statistical model would detect both effects significantly and indicate no interaction term. If interaction is detected, various mechanisms of gene regulations are possible. For example, a detected interaction may indicate that (A) a *cis*-element is present in particular subgenomes of particular lines or that (B, C) a *cis*- and *trans*-factor interact with each other.

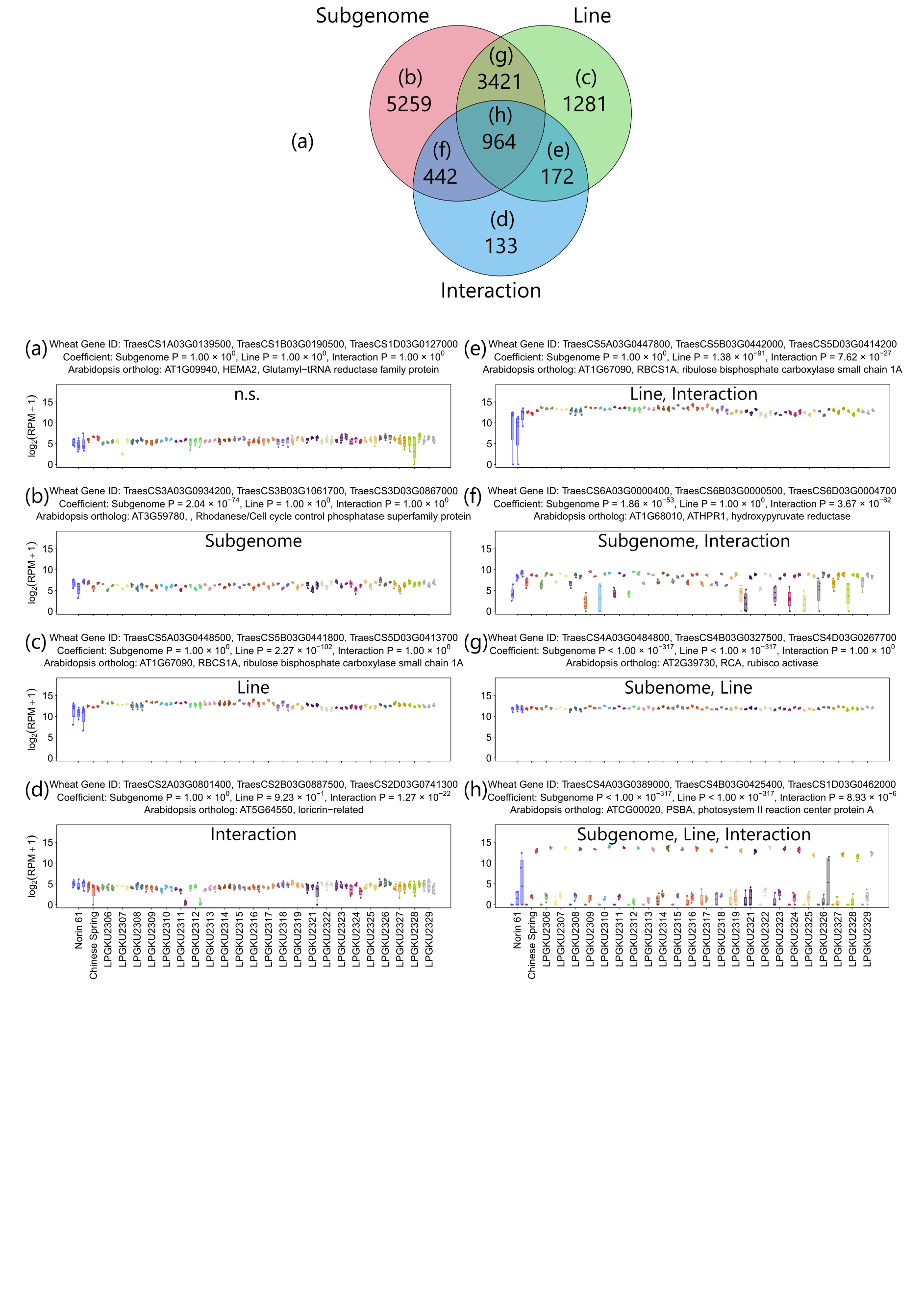

**Supplementary Figure S4 Expression levels of representative genes for which each term is significant in the leaves.** The figure above shows the number of genes where line, subgenome or their interaction term is significant. It also shows the correspondence between the Venn diagram and representative genes; (a) all terms are not significant, (b) only the subgenome term is significant, (c) only the line term is significant, (d) only the interaction term is significant, (e) the line and interaction terms are significant, (f) the subgenome and interaction terms are significant, (g) the subgenome and line terms are significant, and (h) all terms are significant.

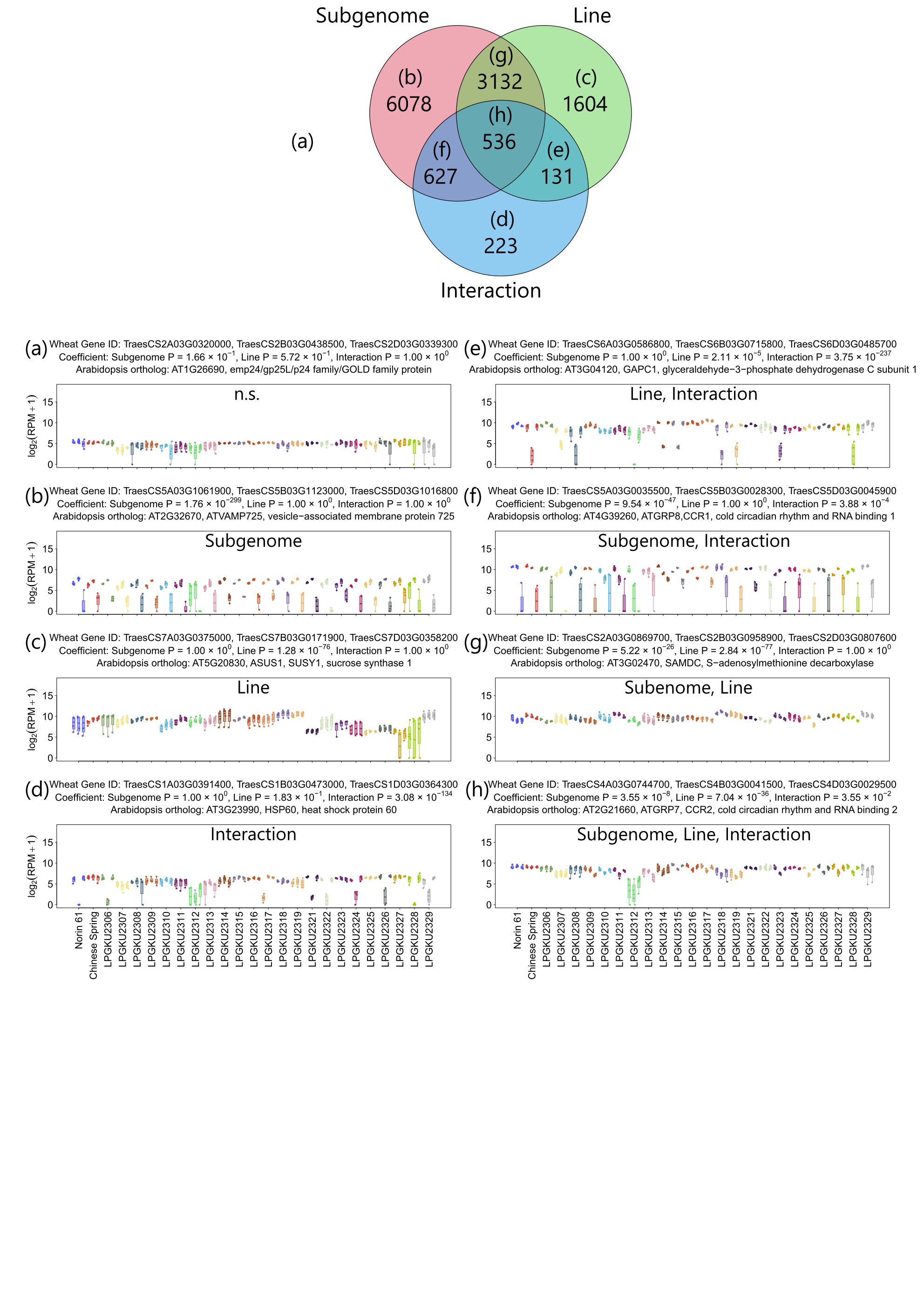

**Supplementary Figure S5 Expression levels of representative genes for which each term is significant in the roots.** The figure above shows the number of genes where line, subgenome or their interaction term is significant. It also shows the correspondence between the Venn diagram and representative genes; (a) all terms are not significant, (b) only the subgenome term is significant, (c) only the line term is significant, (d) only the interaction term is significant, (e) the line and interaction terms are significant, (f) the subgenome and interaction terms are significant, (g) the subgenome and line terms are significant, and (h) all terms are significant.

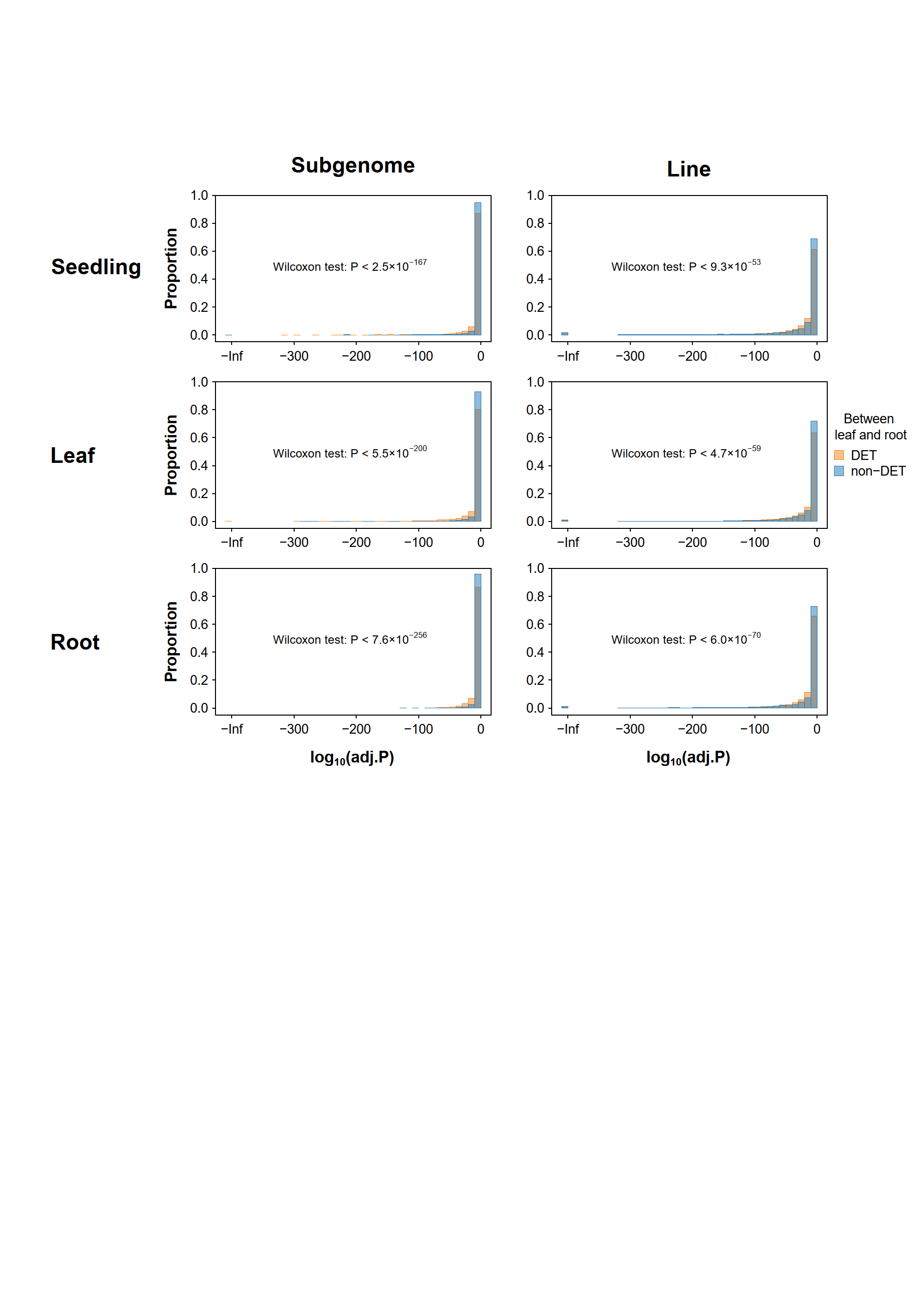

**Supplementary Figure S6 Distribution of adjusted *P*-value in the subgenome and line term.** The x-axis represents log-transformed adjusted *P*-value of the subgenome and line terms in all tissue types as calculated by edgeR. Wilcoxon rank-sum tests were performed to detect the differences in the distribution of adjusted *P*-values between DETs and non-DETs between leaves and roots.
